# Basolateral amygdala parvalbumin interneurons coordinate oscillations to drive reward behaviors

**DOI:** 10.1101/2023.08.09.552674

**Authors:** Kenneth A. Amaya, Eric Teboul, Grant L. Weiss, Pantelis Antonoudiou, Jamie L. Maguire

**Affiliations:** Department of Neuroscience, Tufts University School of Medicine, Boston, Massachusetts, 02111, USA

## Abstract

The basolateral amygdala (BLA) has been implicated in mediating both fear and reward learning. Parvalbumin interneurons (PVs) in the BLA have previously been shown to contribute to BLA oscillatory states integral to fear expression, but whether BLA oscillatory states and PV interneurons also contribute to reward learning is unknown and critical to our understanding of reward processing. Local field potentials in the BLA were collected as animals consumed a sucrose reward, where prominent changes in the beta band (15-30 Hz) emerged with reward experience. Rhythmic optogenetic stimulation of PV interneurons to entrain the BLA network to 20Hz during consumption of one bottle during a two-bottle choice test produced a robust bottle preference. Finally, to demonstrate that PV activity is necessary for reward value use, PVs were chemogenetically targeted and inhibited following outcome devaluation, rendering those animals incapable of using updated reward value information to guide their behavior. Taken together, these experiments provide novel information regarding the physiological signatures of reward learning while highlighting the importance of PV interneurons in reward learning. This work builds upon the field’s established knowledge of PV involvement in fear expression and provides evidence that PV orchestration of unique BLA network states is involved in both learning types.

## Introduction

Use of value information is critical to survival. Prospective considerations of relative outcome values are often used to guide reward seeking and adaptive decision-making, accomplished by using representations of previously experienced rewards. In instrumental conditioning, this is formalized as goal-directed behavior, which is the encoding of the relationship between an action and the outcome it produces ^1^. These behaviors rely on a distributed network of brain areas including the prelimbic area of the frontal cortex, orbitofrontal cortex, mediodorsal thalamus, dorsomedial striatum, and basolateral amygdala (BLA) ^2–7^. Recent advances highlight the BLA as a reward learning hub as it sends and receives projections between these areas ^8^. However, the BLA is also well-documented as a governor of fear behaviors, framing its role as an emotional learning center ^9^.

While the BLA is primarily composed of excitatory principal neurons that project to other emotionally relevant brain areas, microcircuitry in the nucleus includes a complex collection of interneurons that can regulate BLA principal neuron activity. Communication within and between brain regions involved in emotional processing is supported by specific patterns of network activity (oscillations) that have been implicated in the representation, flow, and storage/retrieval of information ^10^, and GABAergic signaling via interneurons has been demonstrated to play a critical role in BLA oscillation generation ^11^. Of particular interest, due to strong evidence supporting their ability to orchestrate oscillations and control fear-related network and behavioral states, are parvalbumin-expressing (PV) interneurons ^12–15^. However, despite the BLA being known to govern both fear and reward behaviors, little is known about reward-related BLA oscillatory activity or whether PVs meaningfully contribute to reward-related network and behavioral states.

To address this, we conducted a series of experiments to dissect PV interneuron contributions to network engagement during reward seeking behaviors. We first characterized BLA oscillations during reward seeking. Then, we demonstrated that optogenetically stimulating PVs to entrain the BLA to the observed reward-related network state is capable of driving reward-seeking behavior. We next examined the necessity of PV interneurons for reward-seeking behaviors by chemogenetically inhibiting them after outcome devaluation, demonstrating a critical role for PV interneurons for performance of both reward seeking and goal-directed instrumental action. Together, these experiments demonstrate that BLA PV interneurons can promote distinct oscillatory states that govern reward-seeking behaviors, providing microcircuit insights into reward processing relevant to disorders such as depression or addiction.

## Materials and Methods

### Experiment 1

#### Animals

Adult C57BL/6J mice (N = 14, 6 male, 8 female, 8-10 weeks old) were housed at Tufts University School of Medicine in a temperature and humidity-controlled environment with a 12-hour light/dark cycle with lights on at 7 AM. All procedures were in accordance with the protocols approved by the Tufts University Institutional Animal Care and Use Committee.

#### Surgery

Mice were anesthetized via intraperitoneal injection of a ketamine/xylazine cocktail (90-120 mg/kg and 5-10 mg/kg, respectively). Before surgery, sustained release buprenorphine (0.5 mg/kg) was administered subcutaneously as post-operative analgesia. For in-vivo local field potential (LFP) recordings, custom fabricated headmounts (Pinnacle Technology, cat. #8201) outfitted with a stainless-steel wire depth electrode (A-M systems, cat. #792300) were inserted unilaterally at the BLA target coordinates: from bregma, -1.5 mm AP, +/-3.3 mm ML, -4.5 mm DV. Headmounts were affixed to the skull using stainless steel screws as ground, reference, and frontal cortex EEG. Animals were allowed to recover for 7 days prior to experimentation.

#### Behavioral apparatus

Animals were trained and tested in 8 identical cylindrical chambers outfitted with two bottles with a pressure-responsive lick spout. Lick event data were collected through a photobeam connected to an Arduino controlled by a python script for lick detection that sent TTL pulses to a Powerlabs LFP acquisition system running LabChart (ADInstruments).

#### Training, testing, LFP recording

Prior to the start of experiments, animals were food restricted to 85% of their *ad libitum* weights. Animals were given access to two bottles, both filled with water, overnight (∼20 hours). Then, after the water-water period, the bottle with fewer licks was changed to contain 20% sucrose and animals were given access for 2 hours. Bottle sides (left vs right) were counterbalanced across animals. LFPs were acquired at 4 kHz and amplified x100 as animals consumed in both the water-water and the water-sucrose phases of the experiment.

#### Analyses and statistical tests

Lick timestamps were collected through an Arduino Due connected to a computer running a custom python script. Lick counts were analyzed by bottle contents (water or sucrose) and average counts were plotted by bottle using R (ggplot2). Lick analyses were performed using generalized linear mixed models to assess the effects of Session Phase, Bottle Type, and their interaction on Lick Counts, with random intercepts for individual animals. Reported statistics include odds ratio estimates, 95% confidence intervals, and p-values for each predictor. A Wilcoxon Signed-Rank Test was used to determine if differences in bottles were present during specific phases, where the V statistic and p-value were included.

Collected lick data were grouped into lick trains when individual licks had an inter-lick interval of less than 3-seconds and were at least 10 seconds in duration. Local field potentials obtained during defined lick-train periods underwent spectral analysis. For assessment of power changes in specific frequency bands over sessions, the mean power following the lick event for each animal was calculated and plotted as the Group mean ±1 SEM by Session. Linear mixed models were used to test fixed effects of Group and Session on Power. A scatterplot and regression fit were used to relate observed Beta power and sucrose preference. Analyses were conducted in R (R Core Team 2016; “stats”, “lme4”, “lmerTest”,”car”). Graphs were generated using R (“ggplot2”) and stylized in Adobe Illustrator.

#### Spectral analysis

Spectral analysis of the LFP/EEG signal was performed using the Short Time Fourier Transform (STFT) with a 5 second Hann window and 50% overlap. The power line noise (59 -61 Hz) was removed and values, filled using nearest interpolation. Outliers in each spectrogram were identified using a two-stage process. Firstly, a time-series was obtained from the mean power across frequencies of each spectrogram. Large deviations, defined as those greater than the mean plus 4 times the standard deviation, were replaced with the median of the non-outlying data. Then, a sliding-window method was applied to detect more subtle outliers based on local context, using 5x the Median Absolute Deviation (MAD) of each 5-minute window. Resulting outliers were removed and replaced with forward fill interpolation of the nearest values. All resulting time-series, obtained from the mean power across frequencies, were manually inspected to remove bad regions that were replaced with median values for each frequency. Any resulting power spectral densities (PSDs) with no apparent peaks were rejected and were not included for further analysis. Each PSD was normalized to its total power across all frequencies analyzed (1 -120 Hz).

#### Histology

Following behavioral experiments, animals were euthanized using isoflurane and tissue was drop-fixed in 4% paraformaldehyde. Following cryoprotection via 30% sucrose saturation, tissue was sectioned, collected, and mounted onto slides with hard-set mounting medium with DAPI for microscope visualization. Images were taken on a Keyence BZ-X700 and estimated depth electrode placements were recorded.

### Experiment 2

#### Animals

Adult PV-Cre mice (B6;129P2-Pvalbtm1(cre)Arbr/J; N = 19, 10 male, 9 female, 9-11 weeks old at surgery) were housed at Tufts University School of Medicine in a temperature and humidity-controlled environment with a 12-hour light/dark cycle with lights on at 7 AM. All procedures were in accordance with protocols approved by the Tufts University IACUC. Animals were genotyped in-house using the primers listed below.

##### Parvalbumin

5’: CATGAGGAGTGGCATACACG

3’: TTCGCGATTTGAGGTCTTCT

#### Surgery

For viral infusions, bilateral burr holes were made in the skull using the BLA target coordinates and mice were bilaterally injected with 250 nL of pAAV9-EF1a-double floxed-hChR2(H134R)-mCherry-WPRE-HGHpa or pAAV9-FLEX-tdTomato using a 33-gauge Hamilton syringe at an infusion rate of 100 nL/min. The syringe was left in place after each infusion for 10 minutes to allow for diffusion of the virus. Two anchor screws were put into the skull and fiber optics were implanted bilaterally (200 um, 0.22 NA; ThorLabs) over BLA and cemented in place with the help of the anchor screws. Animals began the experiment 21 days after surgery to allow for viral expression.

#### Behavioral apparatus

Animals were trained and tested in chambers identical to those described in Experiment 1. Light was delivered through a fiber optic coupled to a blue laser (473 nm, max power = 500 mW, LaserGlow Technologies). The light intensity setting used for these experiments was measured at 8-14 mW from the end of the opto-cannula (ThorLabs, Inc.; 200 um, 0.22 NA). Sine waveforms for photo-stimulation were generated through LabChart as previously described ^12,16^.

#### Training and testing

Animals in Experiment 2 only ever experienced the water-water condition. After a baseline session, one water bottle was paired with optogenetic stimulation of PV interneurons to entrain the network at specific frequencies. Light delivery was triggered by the animal licking the paired bottle and at a frequency of 20 Hz for a duration of 3 seconds, informed by results from Experiment 1. As controls, this process was repeated for 7 Hz and 3 Hz optogenetic stimulations, which has been shown to influence negative valence processing ^12,16^. A tethered baseline session (24 hours) determined which bottle would be paired with light delivery during the test session that immediately followed, lasting another 24 hours. Bottle and light pairings were made based on animal initial bottle preference during their baseline session. For 20 Hz and 7 Hz stimulation sessions, stimulations were paired with the ‘unpreferred’ bottle to evaluate whether stimulating a network state associated with positive valence processing would increase the preference for the ‘unpreferred’ bottle. Conversely, 3 Hz stimulations were paired with the ‘preferred’ bottle to determine whether stimulating a network state associated with negative valence processing would decrease the preference for the ‘preferred’ bottle.

#### Analyses and statistical tests

Lick events were recorded through an Arduino Due connected to a computer running a custom python script. Bottle Preference Scores were calculated as a difference in percentages of consumption on each bottle. Scores were calculated for each animal during each Session and plotted as the mean score with ± SEM, separated by group assignment. This was done for the 20 Hz stimulation pairing and repeated for the 7 Hz and 3 Hz stimulations. Linear mixed models were used to test the fixed effects of Group and Session and their interaction on Preference Score, with random start points for each animal included. Reported statistics include an estimate of each effect, 95% confidence interval, and associated p-value. Analyses were conducted and graphs were created using R (R Core Team 2016; “stats”, “lme4”, “lmerTest”, “car”, “ggplot2”), with Figures being assembled and stylized in Adobe Illustrator.

#### Histology

Following behavioral experiments, animals were euthanized using isoflurane and tissue was drop-fixed in 4% paraformaldehyde. Following cryoprotection via 30% sucrose saturation, tissue was sectioned, collected, and mounted onto slides with hard-set mounting medium with DAPI for microscope visualization. Images were taken on a Keyence BZ-X700.

### Experiment 3

#### Animals

Adult PV-Cre mice (B6;129P2-Pvalbtm1(cre)Arbr/J; N = 18 for the bottle task, 9 male, 9 female; N = 24 for instrumental nose-poke training, 12 male, 12 female; 9 weeks old at surgery) were housed at Tufts University School of Medicine in a temperature and humidity-controlled environment with a 12-hour light/dark cycle with lights on at 7 AM. All procedures were in accordance with protocols approved by the Tufts University IACUC. Animals were genotyped in-house as described in Experiment 2.

#### Surgery

Bilateral burr holes were made in the skull using the BLA target coordinates and mice were bilaterally injected with 250 nL of pAAV9-hSyn-DIO-hM4D(Gi)-mCherry or pAAV9-FLEX-tdTomato using a 33-gauge Hamilton syringe at an infusion rate of 100 nL/min. The syringe was left in place after each infusion for 10 minutes to allow for diffusion of the virus. Incisions were sutured and closed using tissue adhesive. Animals began experiments 21 days after surgery to allow for viral expression.

#### Behavioral apparatus

The apparati used for the Bottle Task in Experiment 3 were identical to those used in Experiments 1 and 2. Flavored 20% sucrose (orange or grape sugar-free Kool-Aid) was used during training and testing. Flavoring was added to 20% sucrose at 2g/L.

Instrumental conditioning was carried out in 4 identical mouse conditioning chambers designed for the 5-choice serial reaction time task housed in sound-attenuating chambers (Med Associates). Each chamber contained a curved rear wall with five 2.5-cm circular holes, 5 cm deep, and 5 cm above floor level. At the entrance of each hole, infrared beam breaks were positioned to detect whether the animal performed a nose-poke response. During training, two of the five nose-poke holes were illuminated with only one of those two producing reward delivery. Each chamber was outfitted with a food area recessed in the center of the front wall (opposite the nose-pokes). Chambers were illuminated by a house light mounted above the floor on the front wall of the chamber. Task events were controlled by computer equipment located adjacent to the chambers.

#### Training and testing

Animals in the Bottle Experiment experienced both the water-water and water-sucrose conditions, with flavored sucrose solutions being used. During Session 1, following the water-water phase, grape flavored sucrose was given. During Session 2, following the water-water phase, orange flavored sucrose was used. Then, during Session 3, animals were given access to both flavors for a 2-hour period to approximate a flavor preference.

During outcome devaluation (Sessions 4-8), animals had their preferred sucrose flavor paired with a nausea-inducing lithium chloride injection (0.15 M, 0.02 mL/g) to form a taste aversion to the flavor. Then, during Session 9, animals began with a brief baseline recording session (5 mins, pre-CNO) before being given injections of clozapine-N-oxide (CNO, 3 mg/kg/mL) or saline. After 30 minutes, the LFP was sampled again during a second recording session (5 mins, during-CNO) before animals were presented with two water bottles filled with flavored water (no sucrose) during a 1-hour choice test. Order of flavor presentation (Sessions 1-2) were counterbalanced across animals. The two baseline recordings (pre and during-CNO, no behavior) were used to characterize BLA network changes as a product of PV interneuron inhibition.

For instrumental conditioning, animals were given one training session each day. No CNO was administered during training. Training began with an initial 30-minute magazine acclimation session where 20% sucrose was delivered at an average rate of 0.1 mL every 45 seconds. Delivery of the sucrose was accompanied by an audible motor raising the dipper arm. The following day, animals were advanced to a fixed-ratio-1 (FR1) schedule of delivery, where one of two illuminated nose-poke ports produced reward delivery. Instrumental training sessions ended after 30 rewards were delivered or 60 minutes had elapsed. Upon completion of three consecutive FR1 sessions (30 rewards earned), animals were advanced to a random-ratio-2 (RR2) schedule of reinforcement, where either 1, 2, or 3 responses were required for reward delivery. After two completed sessions of RR2, animals were advanced to RR3, where 2, 3, or 4 responses were required for reward delivery. After two RR3 sessions, animals were given a brief 5-minute extinction probe to gauge basal levels of instrumental responding and then immediately given one final RR3 training session before being advanced to outcome devaluation and the post-devaluation probe.

Outcome devaluation via satiety was performed by giving animals free access to the sucrose reward (devalued condition) or plain water (non-devalued condition) in the instrumental chambers delivered on a schedule identical to the initial 30-minute magazine training session. Immediately following this, animals were injected with either CNO or vehicle and allowed to rest for at least 30 minutes before behavioral testing. Testing involved a 5-minute extinction probe where nose-poke responses were recorded but no reward was delivered.

#### Analyses and statistical tests

Lick data collected from the Two Bottle Task was analyzed in a manner identical to lick data from Experiment 2, with a Baseline Session and a Post-Devaluation Session (with CNO administered). LFP spectral analysis was performed as described in Experiment 1. Power spectral density plots were generated by normalizing LFP during-CNO administration to pre-CNO baseline activity. Frequency bands were tested using One-Way ANOVAs to test the effect Group assignment on normalized power using specified contrasts to compare control groups and then, separately, PV inhibition to the control animals.

Instrumental nose-pokes and reward magazine entries were recorded through MedPC. Nose-poke rates were calculated as number of events divided by session time. All statistical tests were carried out using R (R Core Team 2016; “stats”, “lme4”, “lmerTest”, “car”), as previously described ^17^. Individualized linear mixed models were used to analyze the effects of dependent variable responding (nose-poke rates) by fixed effects of experimental group and session while accounting for random effects of differences in individual starting values for the dependent variable during Session 1.

#### Histology

Following behavioral experiments, animals were euthanized using isoflurane and tissue was drop-fixed in 4% paraformaldehyde. Following cryoprotection via 30% sucrose saturation, tissue was sectioned, collected, and mounted onto slides with hard-set mounting medium with DAPI for microscope visualization. Images were taken on a Keyence BZ-X700.

## Results

### Basolateral amygdala network dynamics are shaped by reward experience

To evaluate the relationship between reward experience and BLA oscillations, local field potentials (LFPs) were recorded in the BLA during the two-bottle choice test (Fig 1A, 1B). The first session was comprised of a water-water phase, where animals had no preference between the two water bottles (V = 23; p = 0.12). When sucrose was introduced to one bottle, animals preferred that bottle, evidenced by increased lick events revealed by a generalized linear mixed model, showing significant effects of Phase (odds ratio: 0.24; 95% CI: 0.21-0.27; p < 0.001), Bottle Type (OR: 0.05, 95% CI: 0.04-0.06; p < 0.001), and the interaction between Phase and Bottle (OR: 16.83; 95% CI: 14.41 – 19.65; p < 0.001) (Fig 1C). The power spectral density (PSD) was altered during the sucrose consumption period, marked by observable decreases in high-theta power (Fig 1D). To assess the impact of previous experience on network and behavioral states, animals were later given another two-choice session, identical to the first. There, animals again increased the number of licks on the sucrose bottle, where a generalized linear mixed model revealed significant effects of Phase (odds ratio: 0.05; 95% CI: 0.05-0.06; p < 0.001), Bottle Type (OR: 0.00, 95% CI: 0.00-0.01; p < 0.001), and a significant interaction between Session and Bottle Type (OR: 68.71; 95% CI: 59.11 – 79.88; p < 0.001) (Fig 1F). Like the prior lick data, animals licked more during the sucrose phase, driven primarily by licks on the sucrose bottle, and the interaction shows that the number of licks made on each bottle changed over phases. We noticed that in the second session, power in the beta band became more visibly pronounced in the water-normalized sucrose lick PSD (Fig 1G). Based on both the PSDs shown and prior literature that implicates BLA theta activity in emotional learning and beta activity in other reward-encoding areas ^12,13,16,18,19^, we created peri-lick plots of normalized power for those specific frequency bands. Interestingly, those plots show changes in power around the lick event that become more defined in the second session corresponding to increased reward experience (Fig 1E, 1H). To quantify these observations, specifically as a product of experience, the effect of Session was used to assess changes in post-lick normalized power. Interestingly, linear mixed models revealed that beta power increased between sessions (est: 0.23; 95% CI: 0.09 – 0.36; p = 0.002) while high-theta power did not (est: 0.06; 95% CI: -0.04 – 0.16; p = 0.24) (Fig 1I, Fig 1J). To further highlight the importance of beta power with respect to sucrose consumption, bottle preference is significantly predicted by normalized beta power (Fig 1K). A generalized linear mixed model, with family = binomial, was used to model the relationship between Bottle Preference and Beta power, where Preference = 75.18*logPower + 21.21, showing a significant relationship between the two variables. Together, these findings indicate that during reward consumption, beta power is selectively increased as a product of reward experience.

**Figure 1.**
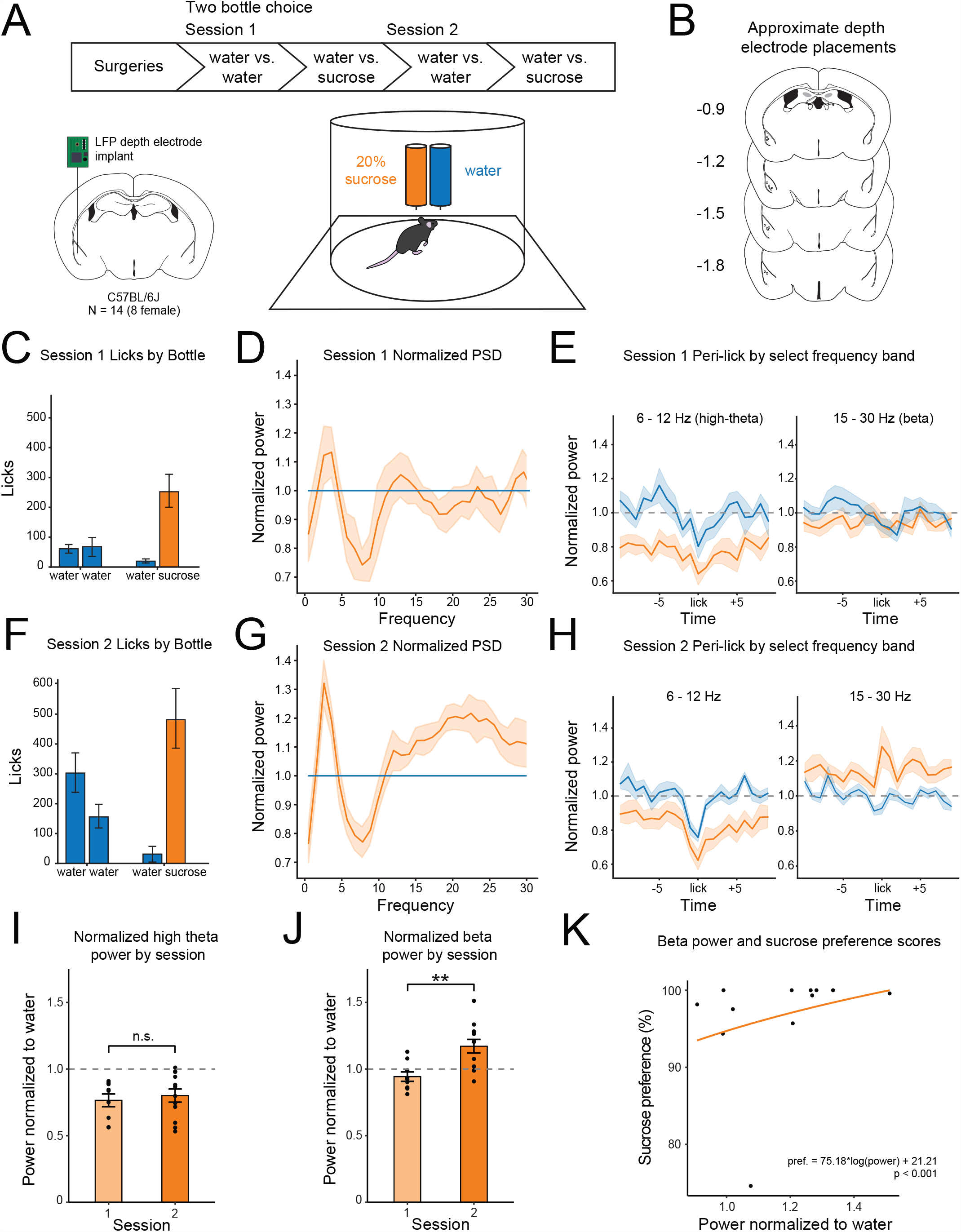
Characterizing reward-related oscillatory states in the basolateral amygdala. A) Schematics diagramming the procedures, showing unilateral LFP depth electrode implant targeting BLA (bottom left), and the two-bottle choice task apparatus, with 20% sucrose and water depicted (sides counterbalanced across animals; bottom right). B) Estimated depth electrode recording locations. C) Lick events by bottle contents during Session 1 of the two-bottle choice task, divided by phase. Left: two identical water bottles presented for 20 hours. Right: one bottle contained 20% sucrose, presented for 2 hours. D) Normalized power spectral density recorded in BLA during lick events of Session 1. E) Peri-lick time plots of select power bands during Session 1. Left: 6-12 Hz (high-theta). Right: 15-30 Hz (beta). F) Lick events by bottle contents during Session 2 of the two-bottle choice task, divided by phase. Left: two identical water bottles presented for 20 hours. Right: one bottle contained 20% sucrose, presented for 2 hours. G) Normalized power spectral density recorded in BLA during lick events of Session 2. H) Peri-lick time plots of select power bands during Session 2. Left: 6-12 Hz (high-theta). Right: 15-30 Hz (beta). I) Normalized high-theta power during sucrose lick events for each Session. Data is selected from the time of lick plus 10 seconds, individual dots represent each animal. Plotted as the mean +/-SEM, no difference between the two sessions. J) Normalized beta power during sucrose lick events for each Session. Data is selected from the time of lick plus 10 seconds, individual dots represent each animal (N = 14). Plotted as the mean +/-SEM, there was a significant increase in beta power from Session 1 to Session 2. K) Logarithmic relationship between beta power and sucrose preference, modeled as Preference = 75.18*log(Norm_Power) + 21.21, where preference is bottle preference from 0-100% and Norm_Power is the normalized beta power observed post-lick.

### PV interneurons can dictate BLA oscillatory states to promote reward seeking

Previous studies have demonstrated the importance of PV interneurons in generating behaviorally-relevant oscillations in the BLA ^12,13^. We tested whether PVs could be similarly leveraged to promote appetitive behaviors. Cre-dependent channelrhodopsin (ChR2) was bilaterally introduced into the BLA of PV-Cre mice. Animals were given access to two identical water bottles in the two-bottle choice task. After a baseline session, a test session followed where one water bottle was paired with rhythmic optogenetic stimulation of BLA PVs (Fig 2A). When pairing one bottle with 20 Hz optogenetic PV stimulations, experimental animals changed their bottle preference to the laser-paired bottle (Fig 2B). A linear mixed model revealed no significant effects of Group (est: 0.83; 95% CI: -0.01 – 1.67; p = 0.051) or Session (est: -0.26; 95% CI: -0.64 – 0.13; p = 0.184) on Bottle Preference Score, but the interaction between Group and Session was significant (est: -0.69; 95% CI: -1.22 – (−0.16); p = 0.012). However, in a later session, when a bottle was paired with 7 Hz optogenetic PV stimulations (a frequency previously associated with safety ^12,16^), there were no behavioral changes observed (Fig 2C). A linear mixed model revealed no significant effects of Group (est: 0.13; 95% CI: -0.81 – 1.07; p = 0.786), Session (est: -0.20; 95% CI: -0.59 – 0.19; p = 0.306), or interaction between Group and Session (est: 0.02; 95% CI: -0.55 – 0.59; p = 0.936) on Bottle Preference Score. Finally, no behavioral changes were observed when the preferred bottle was paired with 3 Hz optogenetic PV stimulations (a frequency associated with fear expression ^12,13,16,18^; Fig 2D). There, a linear mixed model showed no main effects of Group (est: -0.33; 95% CI: -1.24 – 0.58; p = 0.458), Session (est: 0.02; 95% CI: -0.39 – 0.42; p = 0.938), or interaction between Group and Session (est: 0.19; 95% CI: -0.36 – 0.75; p = 0.475) on Bottle Preference Scores. These data suggest that specific 20 Hz entrainment of BLA activity is sufficient to alter reward value use.

**Figure 2.**
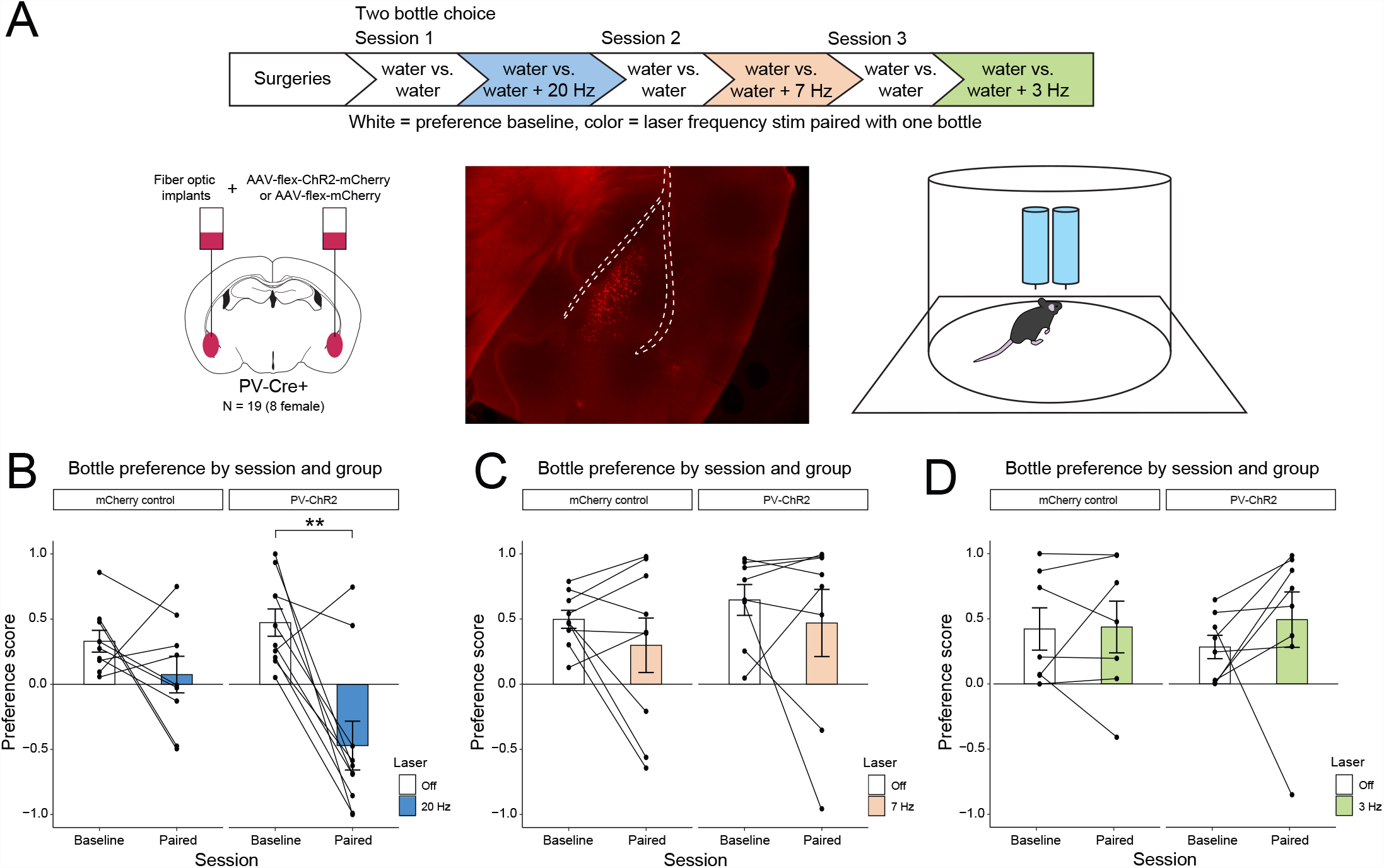
PV-generated oscillatory states selectively promote bottle preference. A) Schematic diagramming procedures (top), showing bilateral viral infusions and fiber optic implant targeting BLA (bottom left), example histology (bottom center) and the two-bottle choice task apparatus, with two water bottles depicted (sides counterbalanced across animals; bottom right). B) Bottle preference by session and group, plotted as a preference score calculated by taking licks on each bottle to assign bottle-specific lick percentages, then subtracting the two values to produce a single preference score. Left: control animals (N = 9) on baseline and during the 20 Hz paired session. Right: experimental animal (N = 10) bottle preferences on the baseline and 20 Hz paired sessions (**p = 0.01). C) Bottle preference by session and group with 7 Hz (high-theta) stimulations (N = 17), no effect of stimulation on preference in either group. D) Bottle preference by session and group with 3 Hz (low-theta) stimulations (N = 15), no effect of stimulation on preference in either group.

### PV interneurons are necessary for BLA-dependent reward behaviors

Next, we sought to establish whether PVs were necessary for reward seeking behaviors known to be BLA-dependent. First, we bilaterally introduced Cre-dependent inhibitory designer receptors exclusively activated by designer drugs (DREADDs) into the BLA of PV-Cre mice before starting animals on the bottle task using two distinctly flavored 20% sucrose solutions (orange and grape, Fig 3A). Animals reduced consumption of the devalued flavored sucrose after outcome devaluation by conditioned taste aversion, shown by a generalized linear mixed model that revealed significant effects of Bottle Identity (OR: 5004.43; 95% CI: 3982.5 – 6288.83; p < 0.001), Session (OR: 30.86; 95% CI: 26.98 – 35.30; p < 0.001), and a significant interaction between Bottle Identity and Session (OR: 0.00; 95% CI: 0.00 – 0.01; p < 0.001) (Fig 3B). To evaluate the role of BLA PVs in using updated value information after devaluation, we chemogenetically inhibited PV interneurons during the flavor recall phase. Inhibition of BLA PVs reduced the power of high theta oscillations in the BLA (mean power +/-SEM: 0.88 +/-0.14) compared to controls (1.43 +/-0.17), t(15) = -2.23, p = 0.042, with no significant effects observed in other frequency bands (see Table 1). During the flavor recall test, animals with PVs inhibited maintained their previously established flavor preference (which was devalued), while controls switched their flavor preference away from the devalued flavor (Fig 3E). This was shown through a linear mixed model where there were significant effects of Group (est: -0.96; 95% CI: -1.38 – (−0.53); p < 0.001) and Session (est: -0.94; 95% CI: -1.13 – (−0.75); p < 0.001) and, importantly, there was a significant interaction between Group and Session, indicating that animals with PV inhibition did not switch bottles after devaluation while control animals did (est: 0.94; 95% CI: 0.67 – 1.21; p <0.001). Between the two control groups, there was not a significant effect of Group (est: 0.14; 95% CI: -0.23 – 0.51; p = 0.44), but there was a significant effect of Session (est: -0.94; 95% CI: -1.13 – (−0.75); p < 0.001), and no significant interaction between Group and Session was observed (est: -0.07; 95% CI: -0.31 – 0.16; p = 0.53). These data demonstrate that BLA PV interneurons are needed to access value changes of specific reward-paired flavors during reward-seeking.

**Table 1.** Effects of PV inhibition across frequency bands during two baseline sessions (pre- and during-CNO) with planned contrasts to compare the two control groups and then compare PV inhibition vs Controls. Table for data from Figure 3E.

| Frequency band | Contrast | t statistic | p-value |
| --- | --- | --- | --- |
| Low theta | Control comparison | 0.465 | 0.65 |
| Low theta | PV vs Controls | -0.741 | 0.47 |
| High theta | Control comparison | 0.283 | 0.78 |
| **High theta** | **PV vs Controls** | **-2.225** | **0.0419** |
| Beta | Control comparison | 0.802 | 0.44 |
| Beta | PV vs Controls | -0.339 | 0.74 |

**Figure 3.**
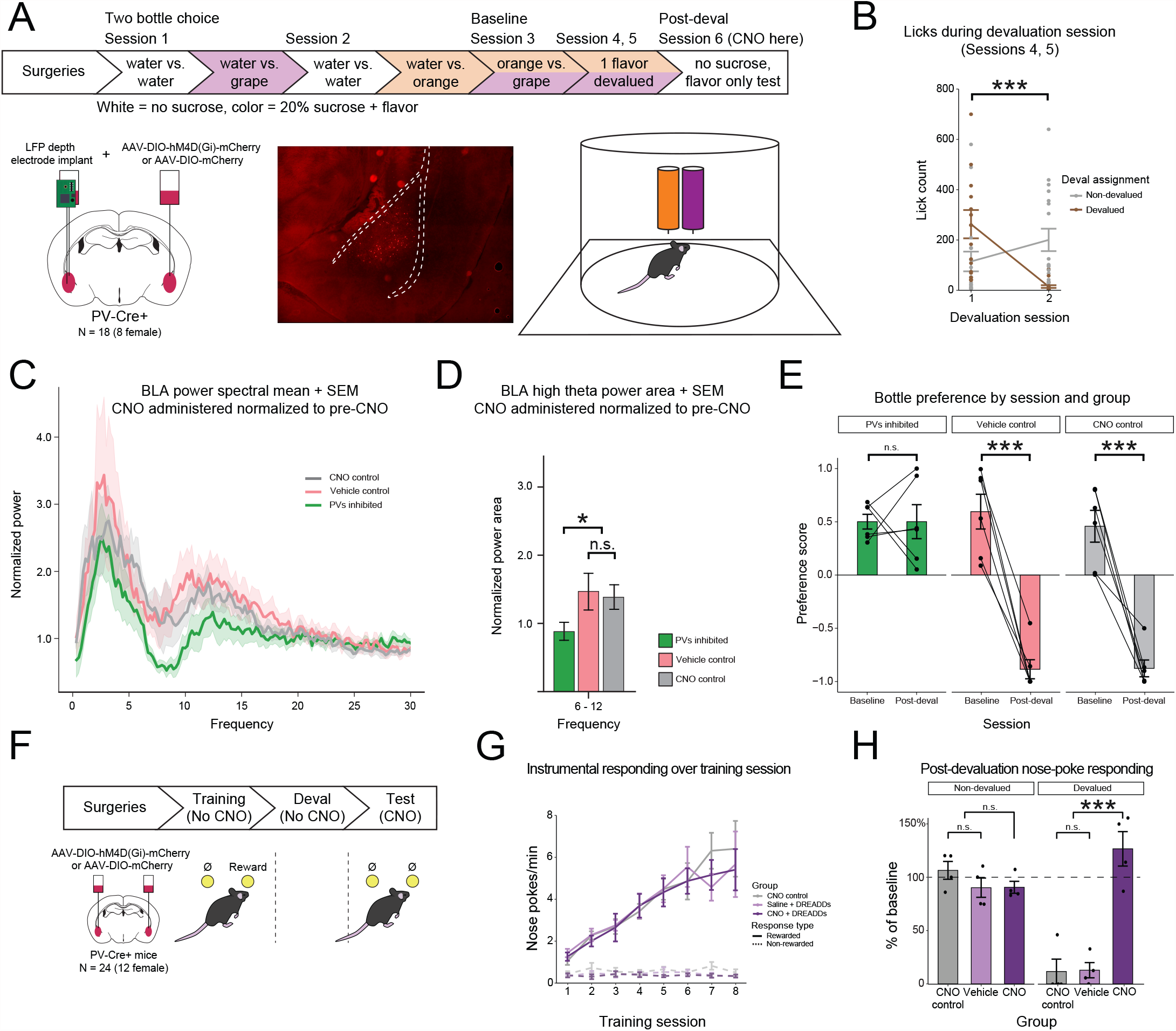
PV activity is required for BLA-dependent reward seeking behaviors. A) Schematic diagramming experimental timeline (top), showing bilateral viral infusions and unilateral depth electrode implant targeting BLA (bottom left), example histology (bottom center), and the two-bottle choice task apparatus, with two flavored sucrose bottles depicted (grape, orange, sides counterbalanced across animals; bottom right). B) Number of licks during devaluation sessions. Session 1 is prior to any devaluation occurring. Animals (N = 18) reduce the number of licks made on the devalued flavor during Session 2. C) Power spectral density (1-30 Hz) recorded in BLA during CNO administration (PVs off-line), normalized to a pre-CNO session of equivalent duration. D) Normalized power in the high-theta band by Group. E) Bottle preference by session and group. Following flavor devaluation, control animals (CNO control, N = 6, and Vehicle control, N = 6) switch their flavor preference. With PVs off-line, animals (N = 6) maintain their preference from their baseline session. F) Schematic diagramming behavioral procedures (top), showing bilateral viral infusions targeting BLA (bottom left), and an instrumental nose-poke task, where responses earned delivery of 0.1 mL 20% sucrose (bottom right). Following training, animals (N = 24) underwent outcome devaluation before finishing experimental testing with reversal training. G) Instrumental training results. All animals learned the task at the same rate and were able to distinguish between the rewarded and non-rewarded manipulanda. H) Post-devaluation responding by group and devaluation assignment. When the outcome was not devalued, all animals (N = 12) continued to perform the task as normal, PV inhibition did not impair instrumental performance. When the outcome was devalued, PV inhibition rendered animals (N = 4) insensitive to the value change, but controls (N = 8) reduced their performance of the action.

Finally, to determine whether BLA PVs are necessary to perform more complex behaviors like instrumental goal-directed behavior, we trained a naive set of animals in an instrumental nose-poke task for delivery of unflavored 20% sucrose outcomes. Cre-dependent inhibitory DREADDs were bilaterally introduced to the BLA of PV-Cre mice before animals began instrumental training (Fig 3F). While no CNO was delivered, all animals learned the task (Fig 3G). A linear mixed model revealed a significant effect of Session on nose-poke rate (est: 0.66; 95% CI: 0.56 – 0.77; p < 0.001) but no significant effect of PV inhibition (est: 0.07; 95% CI: -0.82 – 0.96; p = 0.88) nor an interaction between Group and Session (est: -0.04; 95% CI: -0.19 – 0.10; p = 0.55), indicating that all animals learned to perform the nose-poke action at a similar rate. Following outcome devaluation by satiety, experimental animals, with PVs inhibited, failed to reduce their instrumental responding during the post-devaluation extinction probe (Fig 3H). A two-way ANOVA revealed a significant main effect of Devaluation, F(1,18) = 19.44, p < 0.001, but no main effect of PV inhibition, F(2, 18) = 1.00, p = 0.39. However, there was a significant interaction between Devaluation and PV inhibition, F(2, 18) = 21.57, p < 0.001. These findings show that PV activity is also required to access reward value changes when performing more complex tasks like goal-directed instrumental behavior.

## Discussion

Here, we demonstrate that BLA PV interneurons are critically involved in governing appetitive reward seeking behaviors. We first characterized the relationship between BLA network activity known to be orchestrated by PV interneurons and reward consumption. We then used optogenetics to promote reward seeking through rhythmic PV stimulations at a specific reward-related frequency. Finally, we also demonstrated that flexible behavioral adjustments after reward devaluation requires PV activity. This is true for bottle and flavor preference but also more complex behaviors like instrumental, goal-directed action.

Prior work has associated BLA oscillatory states with fear expression ^12,18^, but BLA activity during reward experience had been less characterized. We demonstrate here that BLA beta oscillations (15 -30 Hz) are of particular importance for reward learning, consistent with previous studies identifying beta oscillations as a marker of learned reward value and behavior in other brain regions ^19,20^. Interestingly, those regions are functionally connected to the BLA, supporting the idea that beta oscillations in the reward network are critical to reward value. Translationally, beta oscillations are a signature of target engagement for some antidepressants ^13^, potentially affecting mood via action on anhedonia, a prominent feature of depression ^21^.

Our PV manipulations indicate that these interneurons are both sufficient and necessary to guide reward-seeking behaviors. As fear expression can be controlled by PV-induced optogenetic BLA entrainment ^12,13,16^, we were similarly able to harness PVs to control BLA activity to promote reward-seeking. Pairing 20 Hz stimulations with licking on one water bottle to create a preference is compelling evidence that BLA activity in the beta band can sufficiently promote appetitive behaviors. This concept is strengthened by our null findings when BLA was entrained to high- and low-theta states which had no effect on reward seeking but have been previously shown to promote safety and fear ^12,13,16,18^. A hallmark study reported invigorated reward-seeking by optogenetically stimulating BLA projections to the nucleus accumbens core at 20 Hz ^22^. Since then, frequency stimulations in the beta range have been commonly used to interrogate neural mechanisms of reward behaviors ^4,23–25^, and our LFP and optogenetics findings add some underlying physiological rationale for those methods. Conversely, inhibiting PVs revealed that they are also necessary for generating reward-related oscillatory states in the BLA. Flavor-specific taste aversion and satiety-induced reduction of goal-directed action are both BLA-dependent behaviors that were impaired by reducing PV activity. Through inhibition, we showed that PVs are necessary for animals to flexibly adjust their behavior after changes in outcome values. This finding aligns with other studies on BLA and reward value ^26^, suggesting that well-documented BLA functions are dependent on PV interneurons.

While these results detail that PVs contribute to governing reward behaviors, mechanisms beyond PV activity related to how downstream communication facilitates behavior are unknown. These include questions like how distinct BLA projection populations are engaged or how reward-network node activity is coordinated to produce behavior. Oscillations offer an exciting avenue to investigate these questions, particularly through the Communication through Coherence theory, where information flow is proposed to be facilitated by rhythmic coherence between neural nodes ^27^. Given that the collective activity of neuron populations in a region are likely the key contributors to local oscillatory states, understanding how specific populations are engaged, would serve the goal of understanding brain function from a systems/network perspective. An intriguing possibility that would require subsequent investigation is that interneurons, potentially PVs given current and prior findings ^12,13,16^, direct information routing through the BLA via selective engagement of specific projection neuron populations.

Presently, PVs appear to be influencing BLA oscillatory states to enable flexible reward pursuit. Through our study, we intended to contribute to the field’s knowledge of reward-related physiology by detailing how learning shapes oscillations while also emphasizing that BLA microcircuitry can govern those observed neural states and associated behaviors. Our findings highlight PVs as key regulators of BLA function as it relates to reward value use, particularly when values change. For disorders of reward processing like in depression or addiction, microcircuit contributions to systems-level deviations in neural processing could provide key mechanistic insights for attempts to swing disordered networks back to healthy states.

